# Synthesizing VERDICT maps from standard DWI data using GANs

**DOI:** 10.1101/2021.02.16.431521

**Authors:** Eleni Chiou, Vanya Valindria, Francesco Giganti, Shonit Punwani, Iasonas Kokkinos, Eleftheria Panagiotaki

## Abstract

VERDICT maps have shown promising results in clinical settings discriminating normal from malignant tissue and identifying specific Gleason grades non-invasively. However, the quantitative estimation of VERDICT maps requires a specific diffusion-weighed imaging (DWI) acquisition. In this study we investigate the feasibility of synthesizing VERDICT maps from standard DWI data from multi-parametric (mp)- MRI by employing conditional generative adversarial networks (GANs). We use data from 67 patients who underwent both standard DWI-MRI and VERDICT MRI and rely on correlation analysis and mean squared error to quantitatively evaluate the quality of the synthetic VERDICT maps. Quantitative results show that the mean values of tumour areas in the synthetic and the real VERDICT maps were strongly correlated while qualitative results indicate that our method can generate realistic VERDICT maps that could supplement mp-MRI assessment for better diagnosis.

## 1 Introduction

Multi-parametric (mp)-MRI, consisting of T2-weighted imaging, diffusion-weighted imaging (DWI) and dynamic contrast enhanced (DCE) imaging, provides non-invasive assessment of the prostate improving the detection and characterization of prostate cancer. However, despite its merits, mp-MRI has some important limitations. In particular, it is characterized by low specificity, provides equivocal findings for around 30% of the patients and correlates moderately with Gleason grade [1].

Towards addressing these limitations, advanced, model-based imaging techniques focus on extracting quantitative metrics that characterize the underlying tissue microstructure in-vivo by modeling the DWI signal [4]. In particular, VERDICT (Vascular, Extracellular and Restricted Diffusion for Cytometry in Tumours) MRI [20,19], which has been recently in clinical trial [14] to supplement the standard mp-MRI for prostate cancer diagnosis, is a model-based, DWI technique that captures the main microstructural properties of cancerous tissue. VERDICT MRI has shown promising results discriminating normal from malignant tissue [19,8,9,25] and identifying specific Gleason grades in-vivo [15,26].

VERDICT MRI combines an optimized DWI acquisition protocol [18] and a mathematical model to estimate microstructural features such as cell size, density, and vascular volume fraction, all of which change in malignancy (Fig. 1). The general model characterizes water diffusion in three primary compartments allowing the estimation of intracellular (fIC), extracellular-extravascular (fEEs) and vascular (fVASC) volume fractions, and cell radius (R). However, the quantitative estimation of these parameters requires a specific DWI acquisition, different to the one widely used for the standard mp-MRI. Specifically, VERDICT MRI requires multiple and higher b-values with different diffusion times (90, 500, 1500, 2000, 3000 s*/*mm^2^) in different directions to derive accurate estimates of the microstructure parameters. Higher b-values improve tumour conspicuity and characterization but require longer scan times. Thus, methods that can estimate the VERDICT maps using standard DWI data from mp-MRI acquisitions would be beneficial for improving diagnostic accuracy without increasing scan time and patient discomfort.

**Fig. 1.**
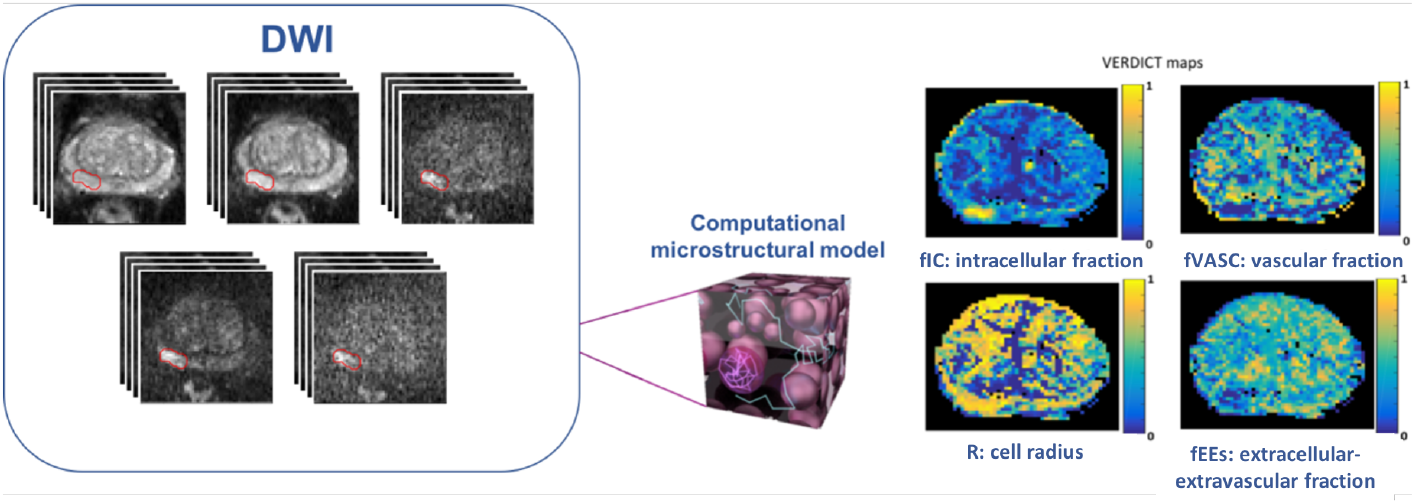
VERDICT MRI framework. It combines an optimized diffusion-weighted imaging (DWI) protocol and a mathematical model to estimate microstructural features of tumours in-vivo.

Recently, several machine learning methods have been proposed to map an input image from one domain to an output image from a different domain aiming to improve the quality of data coming from routine, low-cost acquisitions or scanners or to eliminate the need for multi-modality scanning. Alexander et al. [2] proposed a general framework for image quality enhancement based on patch regression and demonstrated its effectiveness in super-resolution of brain diffusion tensor images and estimation of parametric maps from limited measurements. They further extended this approach with probabilistic deep learning formulation and showed that modeling uncertainty allows for better generalization [23,24]. Oktay et al. [17] proposed an image super-resolution approach based on a residual convolutional neural network (CNN) to reconstruct high resolution 3D volumes from a 2D image enabling more accurate analysis of cardiac morphology. In addition, approaches relying on generative adversarial networks (GANs) [11] have been proposed for super-resolution of structural brain MRI [7,22], endomicroscopy [21] and musculoskeletal MRI [6]. Nie et al. [16] proposed a GAN-based approach to generate CT images from MRI images to eliminate multi-modality scanning. Wolterink et al. [29] used CycleGAN [30] to translate MRI to CT in the absence of paired samples. Wang et al. [27] proposed a semi-supervised approach to synthesize ADC images from T2 images to boost performance of clinical tasks in settings where there is limited supervision. Chiou et al. [10] relied on stochastic translation to translate DWI from mp-MRI to raw diffusion VERDICT MRI to improve segmentation performance.

In this work we also rely on a GAN-based approach [12] to generate VERDICT maps from standard DWI data from clinical mp-MRI acquisitions to obtain microstructural information without requiring a specialized acquisition protocol.

## 2 Methods

### 2.1 Datasets

This study has been performed with local ethics committee approval as part of the INNOVATE clinical trial [14]. The study involved a cohort of 67 men who provided informed written consent.

All participants underwent a standard mp-MRI with a 3.0-T MRI system (Achieva, Philips Healthcare, NL) as part of their standard clinical care. The DWI data was acquired with diffusion-weighted echo-planar imaging sequences. The DWI sequence was acquired with the following imaging parameters: a repetition time msec/echo time msec, 2753/80; field of view, 220×220 mm; section thickness, 5 mm; no intersection gap; acquisition matrix, 168×169mm; b values, 0, 150, 500, 1000, 2000 s/mm^2^. The total imaging time for the clinical diffusion-weighted sequences was 5 minutes 16 seconds.

VERDICT MRI data was acquired with pulsed-gradient spin-echo sequence (PGSE) using an optimised imaging protocol for VERDICT prostate characterization with 5 b-values (90, 500, 1500, 2000, 3000 s/mm^2^) in 3 orthogonal directions [18]. Images with b = 0 s/mm^2^ were also acquired for each b-value. The DWI sequence was acquired with the following imaging parameters: repetition time msec/echo time msec, 2482–3945/50–90; voxel size, 1.25×1.25×5 mm^3^; slice thickness, 5 mm; slices, 14; field of view, 220×220 mm^2^. The images were reconstructed to a 176×176 matrix size. The total imaging time was 12 minutes 25 seconds.

VERDICT MRI maps were generated by using the accelerated microstructure imaging via convex optimization, or AMICO framework [3]. The model has three independent unknown parameters: fIC, R and fEES. fVASC is calculated as fVASC = 1 − fIC − fEES, and the diffusion and pseudo-diffusion coefficients are fixed to dIC = dEES = 2 × 10^−9^m^2^/s, P = 8 × 10^−9^m^2^/s. As in [15], in this study we use the fIC, fEES, fVASC and R.

The regions of interest (ROIs) corresponding to Prostate Imaging Reporting and Data System (PI-RADS) [28] score 3, 4 and 5 were contoured on VERDICT MRI using mp-MRI for guidance by an experienced radiologist reporting more than 2000 prostate MR scans per year.

### 2.2 Proposed Model

Let 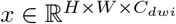, where *H* and *W* are the height and the width of the DWI data and *C*_*dwi*_ = 5, the number of input channels corresponding the different b-values. Let also 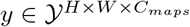, where 𝒴 = [0, 1], *H* and *W* are the height and the width of the VERDICT maps and *C*_*maps*_ = 4, the number of the maps. Our goal is to train a model which takes as input 2D DWI slices (5 b-values) and generates the corresponding VERDICT maps (4 maps).

In this work we use pix2pix framework [12], which has shown great success in natural images, to map standard DWI data from mp-MRI acquisitions to VERDICT maps. As it is illustrated in Figure 2 the framework consists of a generator network *G* and a discriminator network *D*. The discriminator *D* is trained to discriminate between real and synthetic VERDICT maps while the generator *G* maps DWI data to synthetic VERDICT maps (*G* : *x*→*y*) that cannot be distinguished from real VERDICT maps by the discriminator *D*. The adversarial loss, *L*_*GAN*_, can be expressed as

**Fig. 2.**
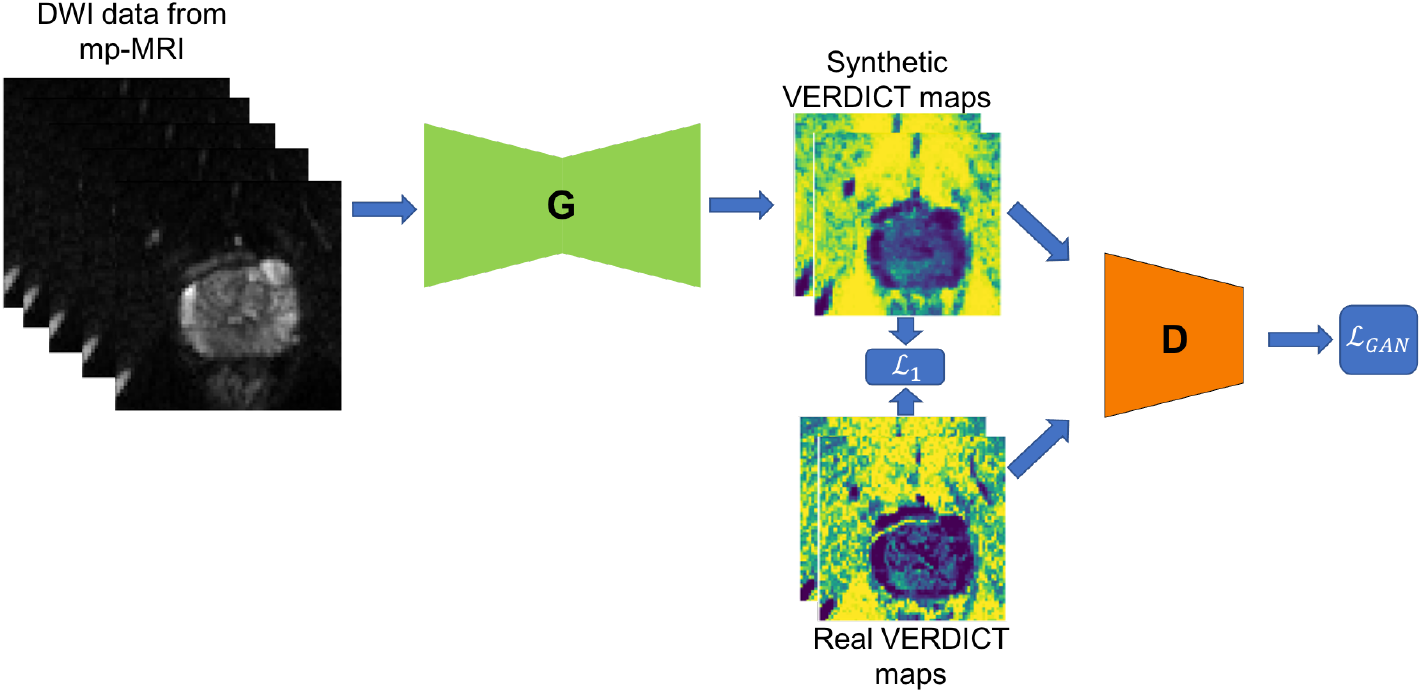
Schematic representation of the proposed framework for synthesizing VERDICT maps form standard DWI data from mp-MRI acquisitions. The discriminator *D* is trained to discriminate between real and synthetic VERDICT maps while the generator *G* maps DWI data to synthetic VERDICT maps (*G* : *x → y*) that cannot be distinguished from real VERDICT maps by the discriminator *D*. A GAN-type loss is used to push the distribution of the synthetic maps closer to the ground truth while the L1 loss ensures that the global and local structure of the synthetic maps do not deviate significantly from the real images.

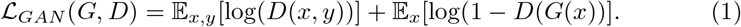

A generator trained solely using the adversarial loss function can synthesize realistic-looking maps which however do not preserve the global and local structure and deviate significantly from the real images. To address this problem and generate maps that both fool the discriminator and are close to the real ground truth maps we use a pixel reconstruction loss, i.e., *L*1 distance. The ℒ_1_ loss can be expressed as

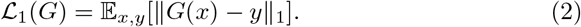

The final training objective can be written as

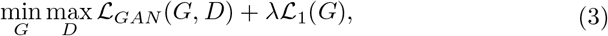

where λ is the weight controlling the importance of the reconstruction loss.

### 2.3 Network architecture

The generator is an encoder-decoder convolutional network based on the U-Net architecture. The encoder consists of 6 convolutional layers followed by batch normalization layers, dropout layers and leaky rectified linear activation units (LeakyRELU). The decoder consists of 6 transposed convolutional layers followed by batch normalization layers, dropout layers and RELU. The last transposed convolutional layer is followed by tanh activation. The output of layer *i* of the encoder is concatenated with the output of the *n*−*i* layer of the decoder, where *n* is the total number of layers, and it is given as input to the next layer of the decoder. The discriminator consists of 3 convolutional layers followed by batch normalization and LeakyRELU. The convolutional layers are 4×4 spatial filters applied with stride 2 and padding 1.

### 2.4 Implementation details

We implement the framework using Pytorch. We train both the generator and discriminator networks using mini-batch stochastic gradient descent and apply the Adam solver with a mini-batch size of 32, and momentum parameters *β*_1_ = 0.5, *β*_2_ = 0.999. We train the networks for 10000 epochs with an initial learning rate at 0.0001 that starts decreasing linearly to 0 after 5000 epochs. We employ dropout as a regularization strategy with dropout rate 50%. We use 60% of the patients for training, 20% for validation and 20% for testing.

## 3 Results

We evaluate the quality of the synthetic maps based on the mean squared error (MSE). Table 1 shows the mean/std of the MSE over 13 test subjects computed on both the entire maps and the prostate region only.

**Table 1.**
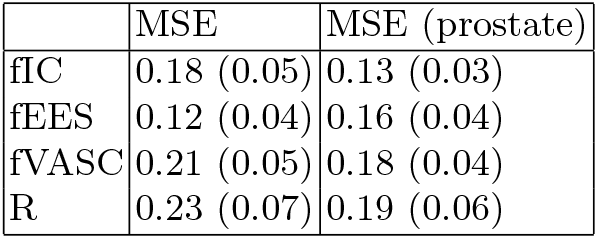
Mean squared error (MSE) calculated on the entire maps and on the prostate region only for the four maps (fIC, fEES, fVASC, R). The results are given in mean (*±*std) format.

|  | MSE | MSE (prostate) |
| --- | --- | --- |
| fIC | 0.18 (0.05) | 0.13 (0.03) |
| fEES | 0.12 (0.04) | 0.16 (0.04) |
| fVASC | 0.21 (0.05) | 0.18 (0.04) |
| R | 0.23 (0.07) | 0.19 (0.06) |

We also calculate the mean value of each ROI on the real and the synthetic maps. Then we compute the correlations between the mean values of the ROIs by computing the Pearson’s correlation coefficient and perform linear regression to quantify the relationships between the mean values in the ROIs. The relationship between the values calculated from the real and synthetic fIC maps is shown in Figure 3 (A). The values show a linear relationship following the regression line fIC_syn_ = 0.87fIC_real_ + 0.09 and the Pearson correlation coefficient is 0.81 (P < 0.05), indicating that there is a strong correlation between the values. Figure 3 (B) shows the relationship between the mean values obtained by the real and the synthetic fEEs maps. The linear relationship is the regression line fEES_syn_ = 0.61fEES_real_ + 0.17 and the Pearson correlation coefficient is 0.74 (P < 0.05). The real and the synthetic fVASC values have a liner relationship given by the regression line fVASC_syn_ = fVASC_real_0.61 + 0.05 and the Pearson correlation coefficient is 0.67 (P < 0.05). The real and the synthetic R values exhibit a linear relationship R_syn_ = 0.53R_real_ + 0.30 and the Pearson correlation coefficient is 0.82 (P < 0.05).

**Fig. 3.**
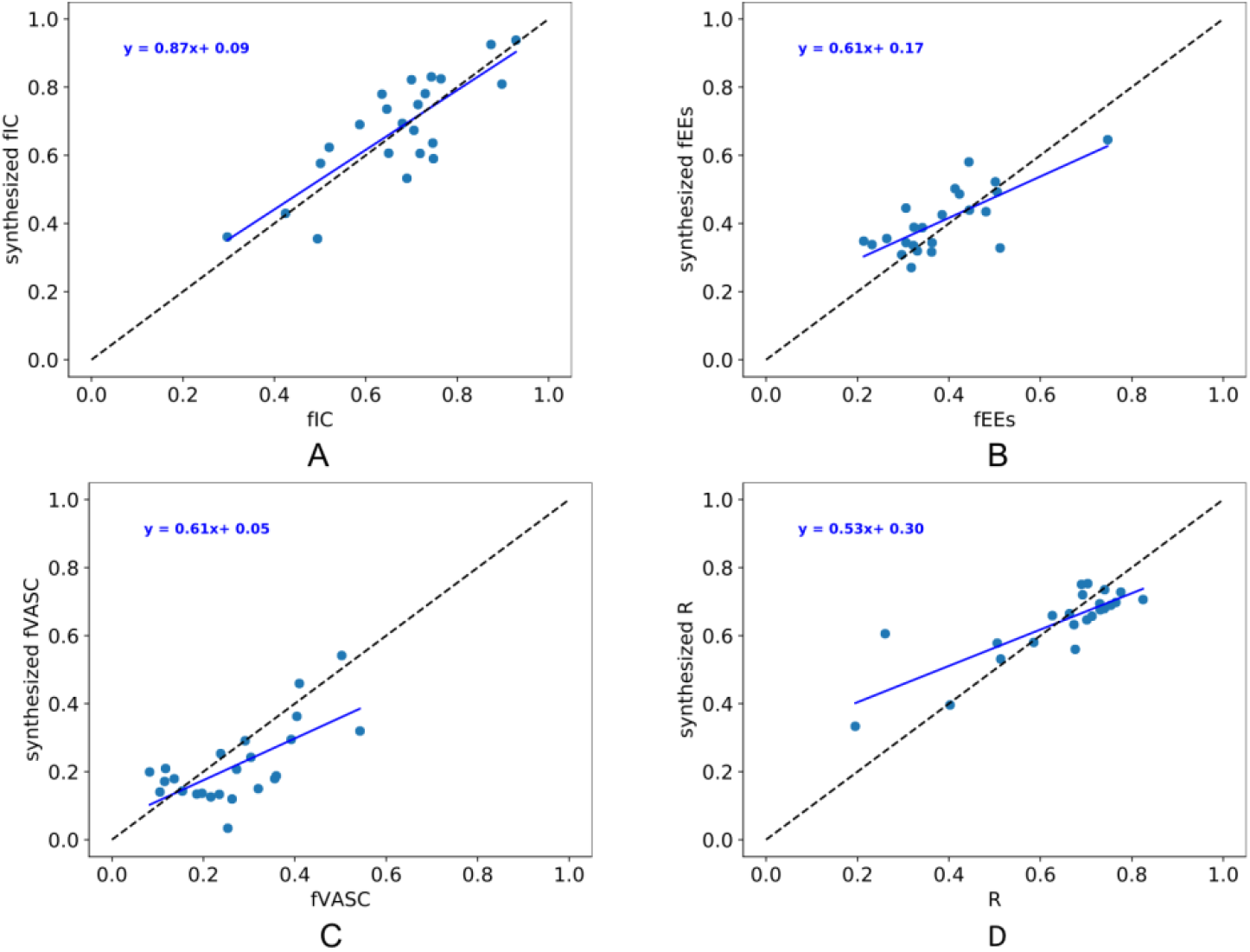
A) Mean values of ROIs calculated from the real fIC as a function of the values calculated from the synthetic fIC. B) Mean values of ROIs calculated from the real fEES as a function of the values calculated from the synthetic fEES. C) Mean values of ROIs calculated from the real fVASC as a function of the values calculated from the synthetic fVASC. D) Mean values of ROIs calculated from the real R as a function of the values calculated from the synthetic R.

Figure 4 demonstrates an example of real and synthetic VERDICT parametric maps for two patients with prostate lesions in the transition zone and the central zone respectively. L1 loss alone leads to reasonable but blurry maps; adding the GAN loss gives much sharper results.

**Fig. 4.**
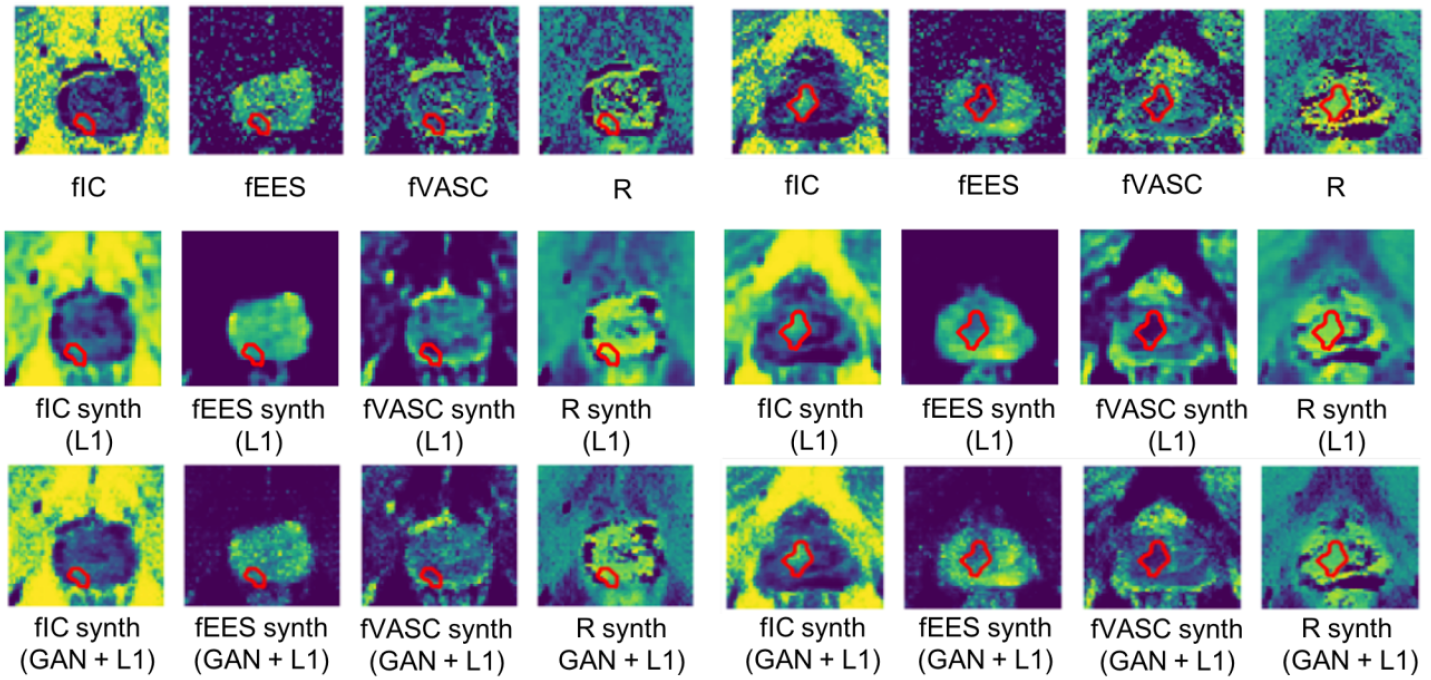
fIC, fEES, fVASC, R maps and the corresponding synthetic maps for two patients with prostate lesions in the transition zone and the central zone respectively. The first row shows the ground truth maps. The second row shows the synthetic maps obtained using the L1 loss alone while the third row shows the maps obtained adding the L1 and GAN losses together.

## 4 Discussion

In this study, we investigate a GAN-based approach for image synthesis for prostate cancer characterization. Our results demonstrate that the proposed approach is viable for generating realistic VERDICT maps from standard DWI data from mp-MRI acquisitions.

Table 1 gives the average MSE calculated between the synthetic and real VERDICT maps. As we can see in the table there is a small difference between the MSE computed using the whole image and using only the prostate region. This indicates that our approach is stable among all regions and all maps, especially the most important one (fIC) that has low error.

In addition the ROI measurements of the synthetic and real VERDICT maps are highly correlated. In particular, there is high correlation for fIC maps (0.81 (P < 0.05)), which have been shown to be the most important in differentiating specific Gleason grades.

Figure 4 shows that the synthetic maps have realistic appearance and preserve important quantitative information. In alignment with the real maps, the synthetic fIC, fEEs, fVASC and R maps clearly depict the lesions which are characterized by high signal in fIC and R maps and low signal in fEES and fVASC maps.

Our objective is to investigate the feasibility of generating VERDICT maps from the clinically-available DWI data from mp-MRI acquisitions to bring the advantages of VERDICT maps in clinical practice. We demonstrate that GAN-based methods have the potential to generate realistic VERDICT maps that preserve important clinical information without requiring specified DWI acquisition protocols, but only using the widely available DWI data from mp-MRI acquisitions. Obtaining VERDICT maps using standard DWI data would reduce acquisition time and patient discomfort. This would also allow us to use already acquired mp-MRI data to get microstructural information.

Despite the good quality of the synthetic maps there are still some limitations. Specifically, the synthetic images are smoother compared to real ones which means that for small ROIs the quantitative values could be wiped out. Methodological improvements that enforce semantic consistency before and after translation [5,13] could resolve this issue and allow the synthesis of high quality VERDICT maps.

## 5 Conclusion

In this work we present an approach for synthesizing realistic VERDICT maps from standard DWI data from prostate mp-MRI acquisitions. Our results indicate that the synthetic maps have realistic appearance and preserve important quantitative information. This could allow the exploitation of VERDICT maps for improved prostate cancer diagnosis without increasing acquisition time and patient discomfort.

